# Enhance Memory CD8^+^ T Cell Formation via Downregulation of Hematopoietic Progenitor Kinase 1 and Sustained Mitochondrial Fitness

**DOI:** 10.1101/2025.04.10.648158

**Authors:** Liuqing Yang, Tantan Wang, Xiangna Guan, Chi He, Hui Shi, Hong Wu, Xuebin Liao

## Abstract

Memory CD8^+^ T cells are pivotal for long-term anti-tumor immunity due to their longevity and rapid response upon encountering tumor cells. Hematopoietic progenitor kinase 1 (HPK1), a member of the Ste20 family of kinases, restricts proximal T cell receptor (TCR) signaling and plays critical roles in T cell priming and activation. However, the impact of HPK1 on CD8^+^ T cell differentiation is not well understood. Here, we demonstrate that chimeric antigen receptor (CAR) T cells derived from patients lacking HPK1 exhibit a stronger memory phenotype. HPK1 deletion promotes the formation of precursor and central memory CD8^+^ T cells, resulting in superior and long-lasting antitumor activity in murine cancer models compared to WT mice. Additionally, HPK1 knockout induces metabolic reprogramming by enhancing oxidative phosphorylation (OXPHOS) and mitochondrial respiration complex activity. Mechanistically, HPK1 deletion downregulates mTOR signaling by reducing the phosphorylation of mTOR and S6. Increased mitophagy activity is observed in HPK1-depleted cells, with upregulation of key mitophagy regulators, including Pink1 and Bnip3l, maintaining mitochondrial fitness. Our study reveals that HPK1 regulates CD8^+^ T cell metabolic reprogramming, guiding cell differentiation and antitumor responses.

## Introduction

Immunotherapy has recently emerged as a revolutionary approach for treating various diseases, particularly cancers^1,2^. The success of immunotherapy hinges on effective CD8^+^ T cell responses, as these cells are the primary effectors in tumor eradication^3^. Memory CD8^+^ T cells, characterized by their longevity, heightened effector functions, and capacity for self-renewal, are pivotal in immunotherapy^4–6^. In advanced melanoma patients receiving anti-CTLA4 therapy, a higher proportion of memory cells of total CD8^+^ T cells in the blood correlates with better efficacy and overall survival^7^. The adoptive transfer of anti-CD19 CAR-T cells containing *ex vivo* expanded memory T cells has also demonstrated significant outcomes^8–12^. The unique properties of memory T cells – including their rapid recall capacity and self-renewing potential – provide a robust biological foundation for developing immunotherapies with sustained clinical benefits against evolving pathogens and heterogeneous tumors.

Metabolic reprogramming is intricately linked to T cell differentiation, activation, and effector responses, with distinct metabolic programs governing the diverse functional states of T cells^13–15^. Effector T cells rely on aerobic glycolysis to meet the biosynthetic demands of proliferating cells, while memory T cells predominantly utilize mitochondrial metabolism, including oxidative phosphorylation (OXPHOS) and fatty acid oxidation (FAO), to sustain long-term survival and effector functions^16^. Inhibition of glycolytic flux – achieved either through limiting glucose uptake^17^ or blocking mitochondrial pyruvate import ^18^– shifts T cells toward oxidative metabolism and enhances memory formation. This transition depends on mitochondrial remodeling – including enhanced fusion, biogenesis, and mitophagy – which collectively maintain the oxidative metabolic state essential for memory cell development and function^19–23^. Additionally, memory T cell differentiation can be enhanced by modulating key metabolic pathways, including mTOR, AKT, and MAPK inhibition or AMPK activation^24–27^. Thus, targeting metabolic pathways to control T cell fate offers a promising strategy for enhancing memory T cell formation and improving immunotherapies.

Hematopoietic Progenitor Kinase 1 (HPK1), a member of the MAP kinase kinase kinase kinase (MAP4K) family, plays a critical role in regulating immune responses^28,29^. HPK1 was initially identified as a negative regulator of T cell receptor (TCR) signaling, inhibiting this pathway by phosphorylating downstream targets like SLP-76 and LAT, attenuating TCR-mediated signaling cascades^30–32^. Further studies have shown that HPK1 broadly influences various aspects of T cell function. HPK1 limits the production of cytokines such as IL-2, IFN-γ, and TNF-α by T cells^33,34^. Increased CD8^+^ tumor-infiltrating lymphocytes are observed in HPK1 catalytic inactive mutant mice^35^. Moreover, HPK1-deficient T cells are resistant to immunosuppressive factors like PGE_2_^35^ and exhibit a less exhausted phenotype^36^. However, the role of HPK1 in regulating memory T cell differentiation remains largely unknown.

In this study, our investigation began with clinical observations of enhanced persistence in HPK1-deficient CD19 CAR-T cell therapies, prompting an in-depth study of their memory T cell properties. Through single-cell transcriptomic profiling and functional validation, we discovered that HPK1 knockout preferentially expands central memory CD8^+^ T cells (CD62L^+^CD27^+^CD8^+^) within the CAR-T population - a subset strongly correlated with sustained remission in patients. Murine models demonstrated that HPK1 depletion promotes memory differentiation and augments antitumor activity. Further mechanistic studies revealed that HPK1 deficiency reprograms T cell metabolism through enhanced oxidative phosphorylation, creating an energy state favorable for memory differentiation. These results establish HPK1 as a key regulator of memory programming in CD8^+^ T cells and provide a biochemical basis for the improved clinical outcomes initially observed.

## Results

### *MAP4K1*^KO^ CD19 CAR-T infusion exhibits more CD62L^+^CD27^+^CD8^+^ T cells

Previous studies have demonstrated that HPK1 knockout (KO) CAR-T cells exhibit significantly enhanced antitumor efficacy and improved safety compared to wild-type (WT) CAR-T cells^36^ However, the impact of HPK1 knockout (KO) on CAR-T cells derived from clinical patient T cells remains unclear. To investigate this, we conducted a study as part of a clinical trial evaluating HPK1^KO^ CD19 CAR-T cells for the treatment of adult acute B-cell lymphoblastic leukemia (NCT04037566). Primary T cells were isolated from five patients and used to generate HPK1^KO^ CD19 CAR-T and WT CD19 CAR-T cells. Single-cell transcriptome sequencing (scRNA-seq) was performed on both groups. After filtering low-quality cells, 80,657 cells were retained, comprising 41,150 HPK1^KO^ and 39,507 WT CAR-T cells. The median number of genes per cell ranged from 3,000 to 4,000 across samples, confirming high-quality data suitable for downstream analysis. To address batch effects, we applied the Harmony algorithm to integrate the datasets. As shown in the figure, cells from different samples are uniformly distributed, indicating successful mitigation of batch effects. (Fig. S1A).

To compare KO and WT CD19 CAR-T cells, we conducted pseudobulk differential expression analysis. The results showed a significant downregulation of *MAP4K1* (the gene encoding HPK1) in KO CAR-T cells, as expected (Fig. 1A). Additionally, genes such as *CD8B* and memory-associated markers, including *CD27* and *SELL* (which encodes CD62L), were upregulated in the KO group (Fig. 1A), suggesting an increased proportion of memory CD8^+^ T cells. We then performed unsupervised clustering of scRNA-seq data and identified 14 cell types based on marker gene expression. These included memory CD8^+^ T cells (T_CD8_memory), effector CD8^+^ T cells (T_CD8_effector), memory CD4^+^ T cells (T_CD4_memory), and cycling CD8^+^ T cells (T_CD8_cyc) (Fig. 1B). To validate our cell type classification, we calculated memory scores using the UCell tool based on memory T cell signature genes (CCR7, TCF7, LEF1, CD62L, CD27, IL7R). T_CD8_memory and T_CD4_memory clusters showed significantly higher memory scores compared to other cell types, confirming their classification (Fig. S1B). A statistical comparison of cell type proportions revealed a significant enrichment of the T_CD8_memory cell type in the KO group (Fig. 1C). Furthermore, the proportion of CD62L^+^CD27^+^CD8+ T cells was markedly higher in KO CAR-T cells compared to WT (Fig. 1D). Using the Single-Cell Entropy (SCENT) tool, we estimated the differentiation potential of CD8^+^ T cells. Differentiation potential scores were significantly elevated in the KO group, indicating that HPK1 knockout enhances the differentiation capacity of CAR-T cells (Fig. 1E).

**Figure 1.**
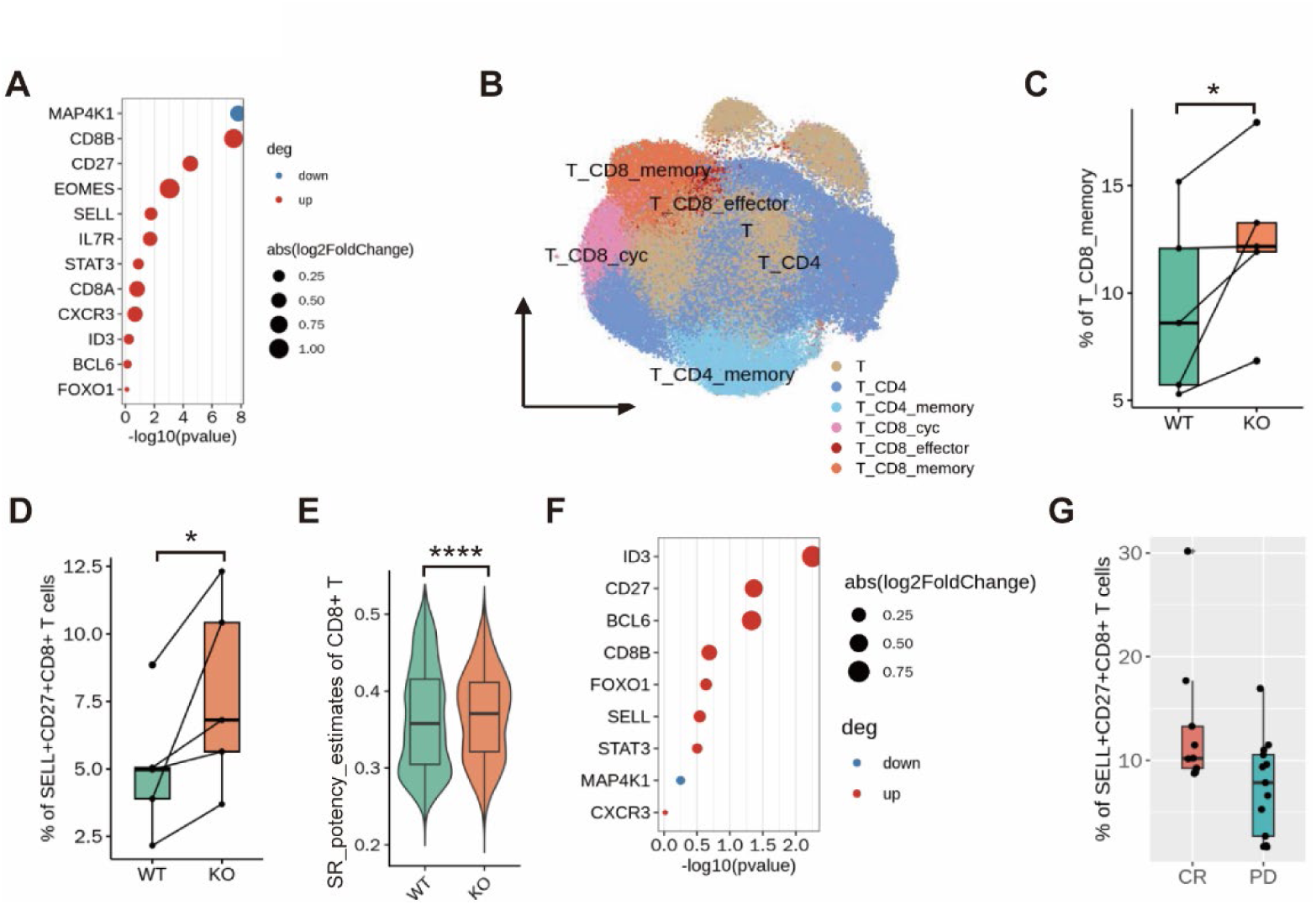
*MAP4K1*^KO^ CAR-T infusion exhibits more CD62L^+^CD27^+^CD8^+^ T cells. A. Upregulated and downregulated genes in HPK1^KO^ CD19 CAR-T cells and WT CD19 CAR-T cells. Red indicates upregulation in the KO group, blue indicates downregulation; B. The cellular composition of CAR-T products was determined by unsupervised clustering of scRNA-seq data, revealing distinct subpopulations; C. The frequency of CD8^+^ memory T cells differed significantly between HPK1-KO and WT CAR-T cells; D. The frequency of CD8^+^ memory T cells (defined as [SELL^+^CD27^+^CD8^+^ T cells]) between HPK-KO and WT CAR-T cells; E. The memory differentiation potential of CD8^+^ T cells in HPK1-KO CAR-T cells and WT controls; F. Upregulated and downregulated genes between CR and PD in dataset of Deng et al. Red indicates upregulation in the CR group, blue indicates downregulation; G. The proportion of CD62L^+^CD27^+^CD8^+^ T cells in CR and PD;

### CD62L^+^CD27^+^CD8^+^ T cells contribute to the clinical efficacy of CAR-T cells

To validate the role of memory CD62L^+^CD27^+^CD8^+^ T cells, we reviewed the literature and reanalyzed published data. In 2020, Deng et al. conducted scRNA-seq on autologous axicabtagene ciloleucel (axi-cel) CD19 CAR-T cell infusion products from 24 patients with large B-cell lymphoma (LBCL) to identify transcriptomic features associated with treatment efficacy and toxicity^37^. The study reported a significantly higher proportion of memory CD8^+^ T cells in CAR-T products from patients with complete remission (CR) compared to those with partial remission (PR) or disease progression (PD). We downloaded the scRNA-seq data from this study and performed pseudobulk differential analysis between the CR and PD groups. The results showed significant upregulation of *CD8B*, *CD27*, and *SELL* in the CR group (Fig. 1F). Focusing on T cell subpopulations, we identified CD4^+^ and CD8^+^ T cells and quantified the proportion of CD62L^+^CD27^+^CD8^+^ T cells (Fig. S1C). The analysis revealed that the proportion of CD62L^+^CD27^+^CD8^+^ T cells was higher in CAR-T cells from patients in the CR group compared to the PD group (Fig. 1G), suggesting that this subset contributes to the clinical efficacy of CAR-T therapy.

To examine the association between HPK1 and the CD62L^+^CD27^+^CD8^+^ T cell subset, we assessed the correlation between HPK1 expression and CD27/CD62L levels across multiple T cell populations in human and murine samples. Using publicly available transcriptomic datasets and published studies, we identified two datasets of CD8^+^ T cells from humans and one from mice for correlation analysis. The results revealed a consistent negative correlation between MAP4K1 expression and levels of SELL (CD62L) and CD27 across all analyzed datasets (Fig. S1D-I). These findings suggest that MAP4K1 functions as a negative regulator of CD62L^+^CD27^+^ memory CD8^+^ T cell differentiation.

### Depletion of HPK1 favors CD8^+^ T cell memory differentiation in mice

To investigate whether the clinical impacts of HPK1 depletion observed in patients also manifest in murine models, we generated mice with CD8^+^ T cell-specific HPK1 depletion by crossing *Map4k1*^fl/fl^ mice with *Cd8*^cre^ mice, resulting in generating T cell-specific HPK1 knockout mice. We then used this type of mice to study CD8^+^ cell differentiation after infection with LCMV Armstrong (LCMV-Arm) (Fig. 2A). On day 8 post-infection, *Map4k1*^−/−^ mice exhibited a significant increase in memory precursor cells (MP; CD8^+^KLRG1^−^CD127^+^) in both spleen and lymph nodes compared to *Map4k1*^fl/fl^ mice (Fig. 2B; Fig. 2A). Furthermore, the percentages of CD62L^+^CD27^+^ memory cells and central memory cells (CD44^+^CD62L^+^CD27^+^) were more than twofold in *Map4k1*^−/−^ mice as compared to *Map4k1*^fl/fl^ mice (Fig. 2C-D; Fig. S2B-C). These phenotypic changes were sustained at days 20 and 30 post-LCMV infection (Fig. 2C-D; Fig. 2B-C), consistent with a *Map4k1*-dependent shift in CD8^+^ T cell fate commitment toward memory differentiation.

**Figure 2.**
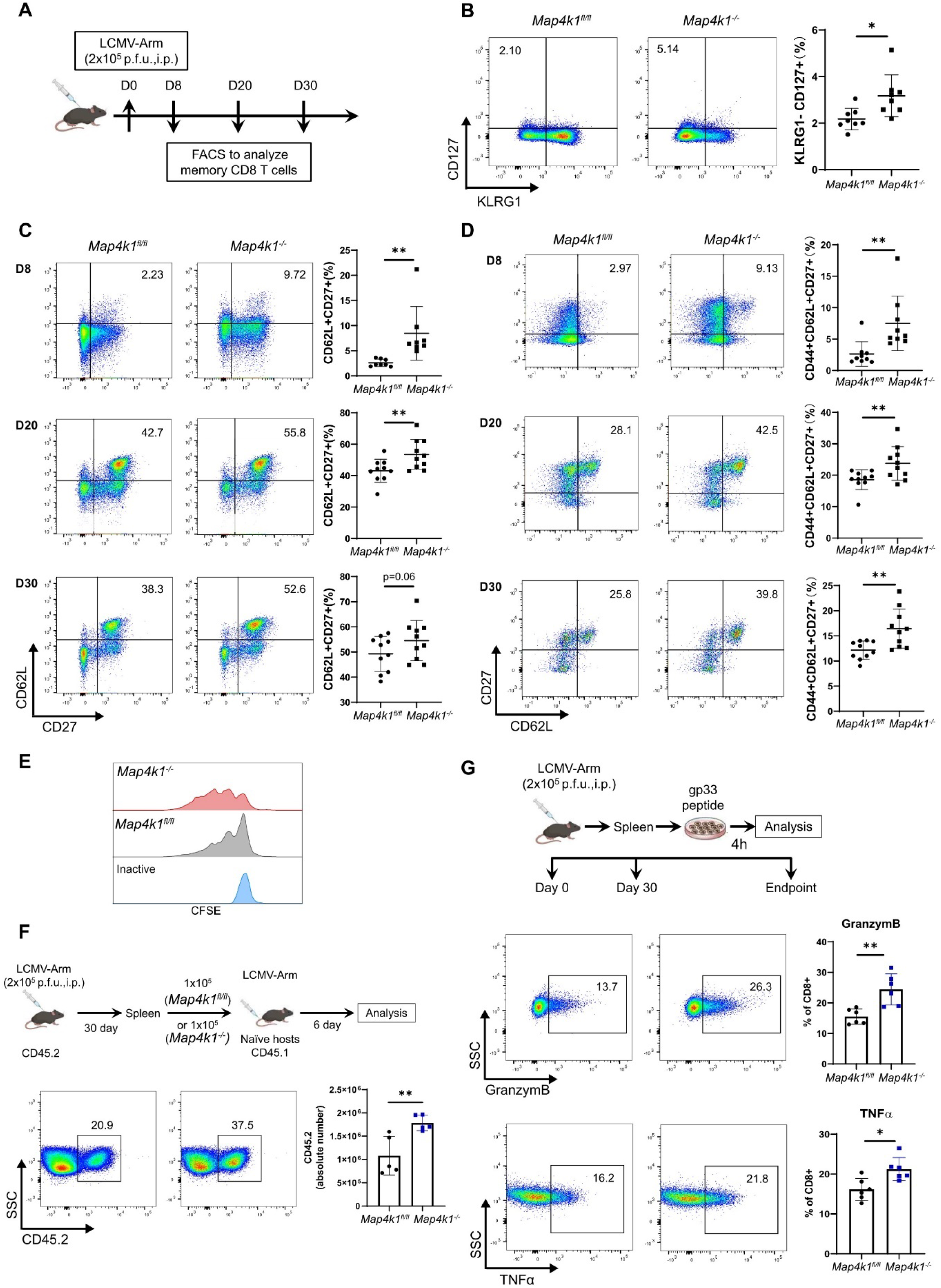
HPK1-defect mice favor memory CD8^+^ T cell differentiation. A. Diagram of LCMV-Armstrong infection model using *Map4k1*^fl/fl^-*CD8*^Cre^ mice (*Map4k1*^−/−^ below) and *Map4k1*^fl/fl^ mice as control (*Map4k1* ^fl/fl^ below). B. Flow cytometry (left) and quantification of frequency (right) of CD127^+^KLRG1^−^ cells in total CD8^+^ T cells from spleen at day 8 post-infection. C-D. Flow cytometry (left) and quantification of frequency (right) of CD62L^+^CD27^+^ cells (C) and CD44^+^CD62L^+^CD27^+^ cells (D) in total CD8^+^ T cells derived from the spleen at indicated times post-infection. E. Cell proliferation of CD8+ T cells separated from LCMV-infected mice upon stimulation *in vitro*. F. Diagram of the LCMV-recall assay (upper), flow cytometry (lower left), and quantification of the number (lower right) of donor-derived CD8^+^ cells following LCMV-Arm secondary challenge. G. Schematic of secondary stimulation of CD8^+^ T cells from LCMV-infected mice *in vitro* (upper). The intracellular expression of Granzyme B (middle) and TNFα (bottom) are represented by flow cytometry plots (left) and quantification of frequency (right).

Memory cells are characterized by their robust proliferative response upon antigen rechallenge. Thus, we assessed whether *Map4k1* depletion enhances the proliferative potential of memory CD8^+^ T cells *ex vivo*. Splenic CD8^+^ T cells isolated from both *Map4k1*^fl/fl^ and *Map4k1*^−/−^ mice at day 30 post-infection were labeled with CFSE and stimulated with LCMV-Arm gp33 peptides. After 4 days, *Map4k1*^−/−^ CD8^+^ T cells exhibited significantly superior proliferation compared to *Map4k1*^fl/fl^ CD8^+^ T cells (Fig. 2E), suggesting enhanced capacity for expansion and renewal of the memory cell pool.

To further investigate this, we adoptively transferred CD45.2-labeled *Map4k1*^−/−^ CD8^+^ T cells isolated from mice on day 30 after LCMV infection into CD45.1 mice, followed by LCMV rechallenge (Fig. 2F). As anticipated, *Map4k1*^−/−^ CD8^+^ T cells exhibited significantly enhanced proliferative expansion upon antigen rechallenge relative to *Map4k1*^fl/fl^ controls (nearly double the expansion; Fig. 2F).

Additionally, we examined the cytotoxic function of memory CD8^+^ T cells upon rechallenge. CD8^+^ T cells isolated from the spleens of both *Map4k1*^fl/fl^ and *Map4k1*^−/−^ mice on day 30 after LCMV infection were stimulated with gp33 peptides *in vitro.* After 4 hours, *Map4k1*^−/−^ CD8^+^ T cells consistently showed increased production of cytotoxic function-related cytokines (TNFα and granzyme B) (Fig. 2G). Collectively, these data indicate that *Map4k1* deficiency enhances the generation of functional memory CD8+ T cells, as evidenced by their heightened proliferative capacity and augmented effector functions.

### HPK1 deficiency in CD8^+^ T cells enhances their long-term antitumor functionality

As a chronic disease, cancer requires sustained T cell-mediated immunity, underscoring the critical importance of memory T cell responses in effective immunotherapy ^6^. Based on prior findings in infection models, we postulated that HPK1 may regulate memory CD8^+^ T cell differentiation within the tumor microenvironment. To test this hypothesis, we compared tumor growth kinetics in *Map4k1*^−/−^ versus *Map4k1*^fl/fl^ mice. (Fig. 3A). In a syngeneic murine B16-OVA melanoma model, *Map4k1* depletion markedly suppressed tumor growth compared to *Map4k1*^fl/fl^ mice (Fig. 3B). Subsequently, we analyzed CD8^+^ T cell subsets in the spleen, tumor-draining lymph nodes, and blood of B16-OVA-bearing mice on day 15. *Map4k1*^−/−^ mice exhibited significantly higher frequencies of CD62L^+^CD27^+^ memory cells across all tissues compared to *Map4k1*^fl/fl^ mice (Fig. 3C; Fig. S3A), underscoring HPK1’s role in regulating memory T cell differentiation within the tumor microenvironment.

**Figure 3.**
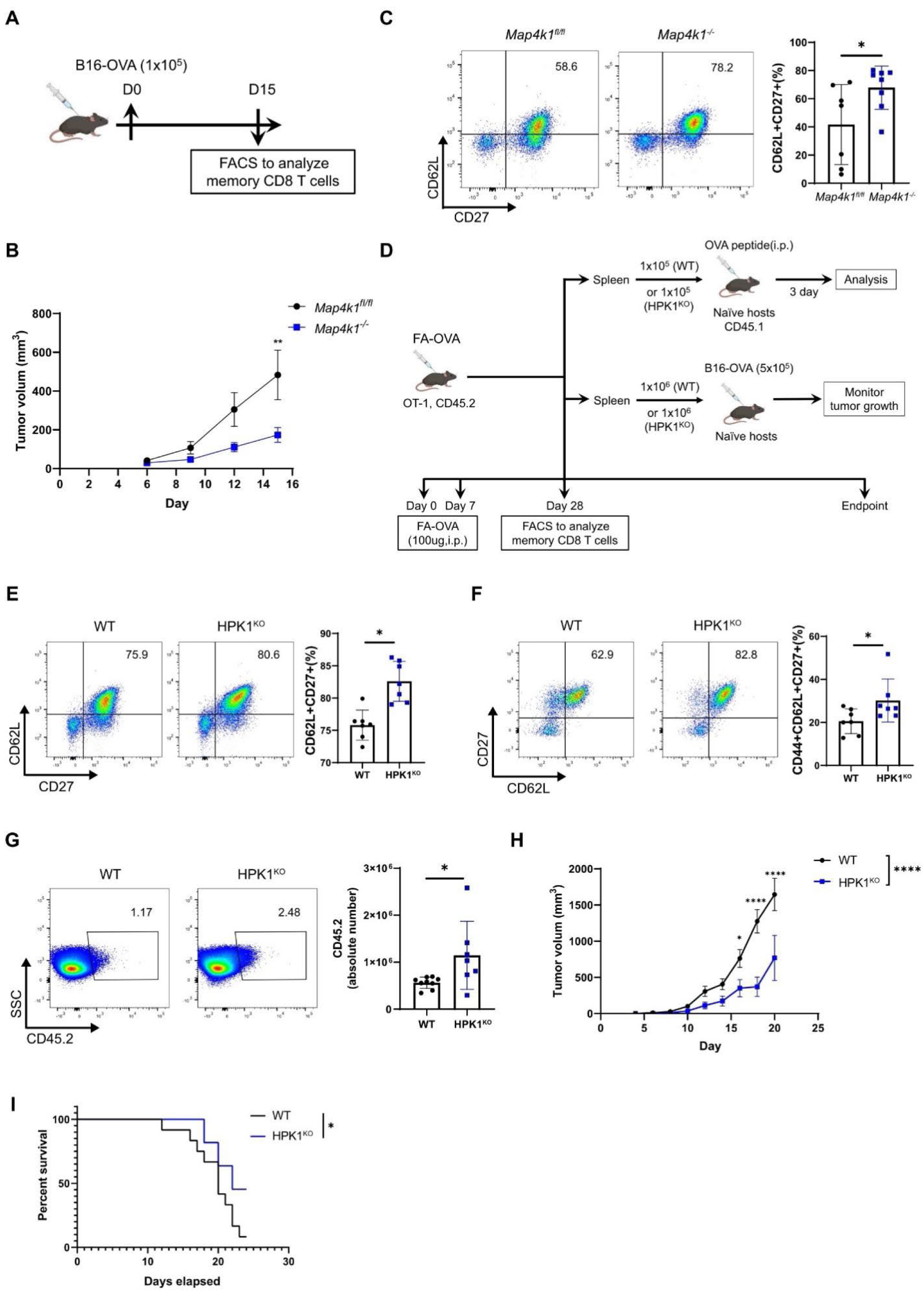
Depletion of HPK1 in CD8^+^ cells improves long-term antitumor capacity. A-C. B16-OVA tumor modal using *Map4k1*^fl/fl^-*CD8*^Cre^ mice (*Map4k1*^−/−^) and *Map4k1*^fl/fl^ mice as control (*Map4k1* ^fl/fl^). (A). Experimental diagram. (B). Tumor growth curve. (C). FACS analyses of CD62L and CD27 expression in splenic CD8^+^ T cells at day 15. D. Diagram of OVA vaccination model and recall assays. E-F. Flow cytometry (left) and quantification of frequency (right) of CD62L^+^CD27^+^ cells (E) and CD44^+^CD62L^+^CD27^+^ cells (F) from total splenic CD8^+^ T cells on day 28 post-OVA vaccination. G. Flow cytometry (left) and quantification (right) of donor-derived CD8^+^ cells after OVA peptide re-challenge. H-I. Tumor growth curve (H) and survival curve (I) of B16-OVA tumor-bearing mice in (D).

To further explore HPK1’s impact on antigen-specific CD8^+^ T cell responses, we crossed HPK1 knockout mice with OT-I mice. Both naïve OT-I (referred to as WT) and HPK1 knockout OT-I (referred to as HPK1^KO^) mice were intraperitoneally immunized with OVA peptides (Fig. 3D). By day 28 post-vaccination, HPK1^KO^ mice demonstrated significantly elevated frequencies of splenic CD62L^+^CD27^+^ memory T cells and CD44^+^CD62L^+^CD27^+^ central memory T cells compared to wild-type controls (Fig. 3E-F; Fig. 3B-C). These findings implicate HPK1 in the regulation of antigen-specific memory T cell development.

To assess the proliferative potential of memory T cells, we conducted adoptive transfer experiments. Splenic CD8^+^ T cells from vaccinated WT and HPK1-deficient mice (CD45.2) were transferred into naïve WT recipients (CD45.1), followed by OVA peptide injection (Fig. 3D). Notably, HPK1^KO^ T cells exhibited enhanced expansion in the spleen upon antigen rechallenge compared to WT T cells, indicating superior recall memory function in the absence of HPK1 (Fig. 3G; Fig. S3D). These results establish HPK1 as a key regulator of memory T cell differentiation and highlight its therapeutic potential as a target for improving vaccine-induced immune memory.

Additionally, splenic CD8^+^ T cells from WT and HPK1^KO^ mice (CD45.2) on day 28 after post-vaccination were adoptively transferred into naïve WT recipients (CD45.1), followed by subcutaneous inoculation of B16-OVA tumor cells (Fig. 3D). HPK1^KO^ recipients exhibited significantly reduced B16-OVA tumor growth compared to WT controls (Fig. 3H), accompanied by markedly improved survival rates (Fig. 3I). In summary, HPK1 deficiency promotes CD8^+^ memory T cell differentiation, sustaining durable antitumor immunity. These findings provide a mechanistic basis for targeting HPK1 to enhance cancer immunotherapy.

### HPK1 inhibition of CD8^+^ T cell induces metabolic reprogramming

To define the molecular mechanisms underlying *Map4k1*-dependent CD8^+^ T cell differentiation, we performed comparative RNA sequencing of sorted CD8^+^ T cells from *Map4k1*^−/−^ and *Map4k1*^fl/fl^ mice. This analysis revealed significant alterations in the transcriptional program governing T cell memory formation (Fig. S4A). *Map4k1* deletion significantly upregulated genes associated with T cell development, such as *Tcf7*, *Id3*, and *Stat3* (Fig. 4A), crucial for orchestrating the transition from naïve to memory T cells^38^. Gene Set Enrichment Analysis (GSEA) confirmed a distinct enrichment of genes related to memory CD8^+^ T cells in *Map4k1*^−/−^ CD8^+^ T cells (Fig. 4B; Fig. S4B). Conversely, genes associated with T cell exhaustion, including *Tbx21*, *Ctla4*, *Id2*, and *Stat4*, were downregulated upon *Map4k1* deletion (Fig. 4A), suggesting a mitigation of exhaustion processes and prolonged CD8^+^ T cell responsiveness.

**Figure 4.**
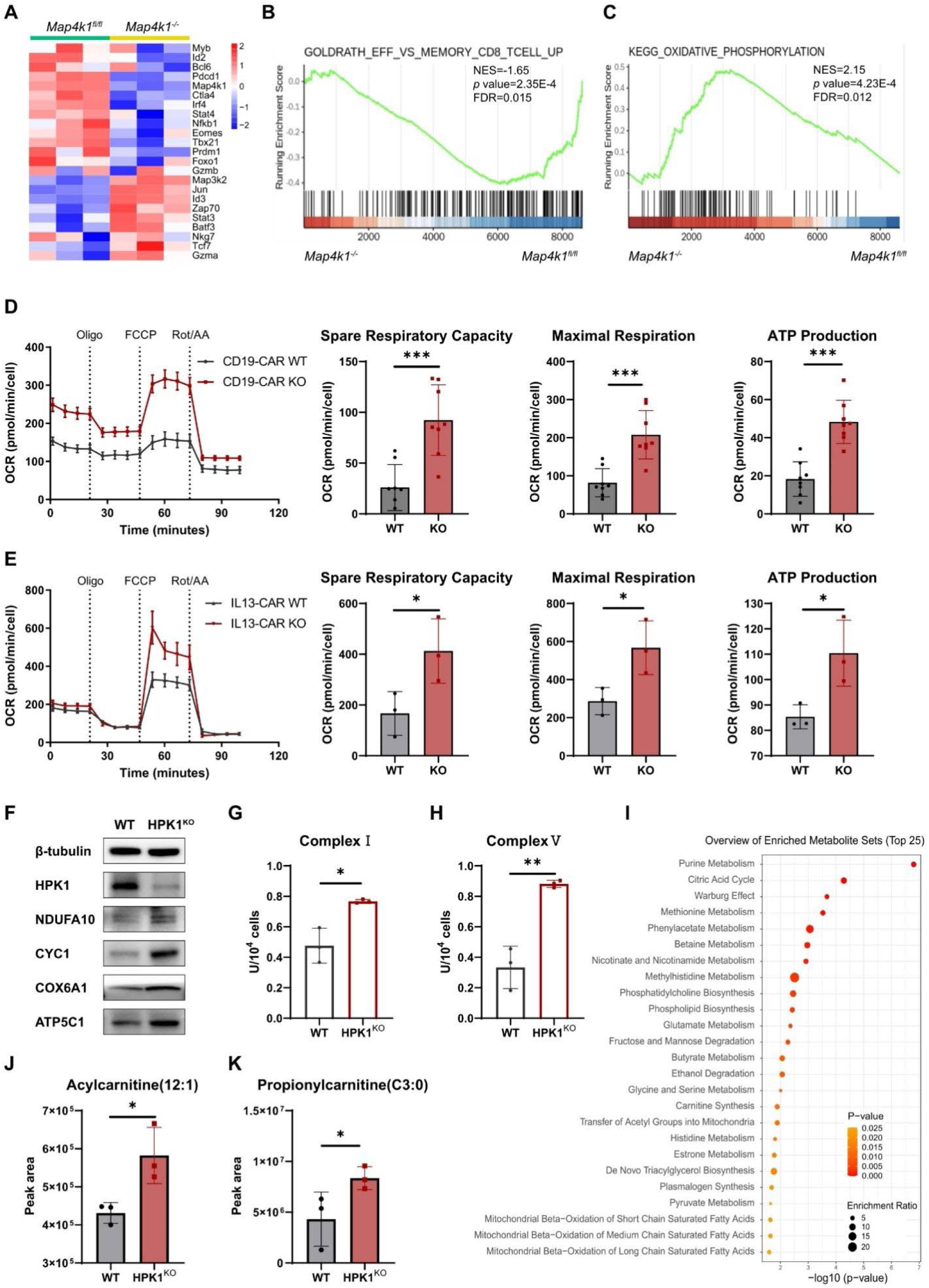
HPK1 influences CD8^+^ T cell metabolic reprogramming. A. Heat map of chosen differentially expressed genes between CD8^+^ T cells from *Map4k1*^fl/fl^ and *Map4k1*^−/−^ mice. B-C. GSEA of effector versus memory enrichment (B) and the gene set related to oxidative phosphorylation (C) in comparisons between CD8^+^ T cells from *Map4k1*^−/−^ and *Map4k1*^fl/fl^ mice. D-E. OCR of WT and HPK1^KO^ CD19-CAR T cells (D) and IL13-CAR T cells (E) measured by Seahorse assay (left). Quantification of spare respiration capacity, maximal respiration, and ATP production of indicated cells (right). Oligo, oligomycin; FCCP, carbonyl cyanide-p-trifluoromethoxyphenylhydrazon; Rot/AA, rotenone and antimycin A. F. Immunoblot of indicated protein of CD8^+^ T cells from WT and HPK1^KO^ mice. G-H. Quantification of activity of mitochondrial complex I (G) and complex V (H) of CD8^+^ T cells from WT and HPK1^KO^ mice. I. Pathway enrichment of metabolites from CD8^+^ T cells derived from WT and HPK1^KO^ mice. J-K. Quantification of acylcarnitine (J) and propionylcarnitine (K) by LC-MS in CD8^+^ T cells from WT and HPK1^KO^ mice.

Further pathway analysis demonstrated that *Map4k1* deletion significantly alters metabolic gene expression, underscoring HPK1’s role in regulating metabolic reprogramming. Consistent with the central role of metabolic pathways in controlling T cell differentiation and effector function^39^, these alterations were particularly pronounced in mitochondrial energy metabolism. Specifically, HPK1-deficient CD8^+^ T cells displayed marked upregulation of oxidative phosphorylation (OXPHOS) and electron transport chain (ETC) genes (Fig. 4C; Fig. S4C), indicating that HPK1 normally restricts these pathways. This shift toward enhanced mitochondrial respiration suggests HPK1 constrains metabolic fitness in CD8^+^ T cells, potentially impacting their cytotoxic capacity and long-term survival.

To investigate the metabolic consequences of HPK1 deletion in CD8^+^ T cells, we performed mitochondrial functional assays using extracellular flux analysis (Seahorse assays). HPK1-deficient (HPK1^KO^) CD8^+^ T cells demonstrated a significantly elevated oxygen consumption rate (OCR) compared to wild-type (WT) controls (Fig. S4D), indicative of enhanced oxidative phosphorylation. This metabolic shift suggests that HPK1 constrains mitochondrial respiration in CD8^+^ T cells, thereby potentially augmenting their capacity to sustain effector responses and support memory differentiation. Furthermore, HPK1-deficient CD8^+^ T cells exhibited significantly increased spare respiratory capacity, maximal respiration, and ATP production (Fig. S4D), demonstrating enhanced mitochondrial metabolic capacity. These results establish HPK1 as a critical negative regulator of mitochondrial bioenergetics in CD8^+^ T cells.

Given the critical role of T cell dysfunction in limiting CAR-T therapy efficacy, we investigated HPK1’s metabolic regulation in human CAR-T cells. Seahorse metabolic profiling demonstrated that HPK1 deficiency enhances mitochondrial fitness, evidenced by significantly increased oxygen consumption rate (OCR) and spare respiratory capacity (Fig. 4D-E). This conserved metabolic phenotype - observed in both murine memory T cells and human CAR-T cells - positions HPK1 inhibition as a dual strategy to simultaneously enhance memory differentiation while preventing exhaustion-associated metabolic collapse, thereby potentially improving CAR-T cell persistence and therapeutic outcomes.

To mechanistically link HPK1 deficiency to enhanced oxidative metabolism, we further analyzed mitochondrial respiratory chain components at molecular and functional levels. Western blotting demonstrated increased protein expression of key respiratory subunits (NDUFA10 [complex I], CYC1 [complex III], COX6A1 [complex IV], and ATP5C1 [complex V]) in HPK1^KO^ CD8^+^ T cells (Fig. 4F), while qPCR confirmed upregulation of corresponding mitochondrial genes (Fig. S4E). Importantly, enzymatic assays revealed significantly elevated activities of complexes I and V (Fig. 4G-H), providing direct functional evidence that aligns with the enhanced oxidative metabolism observed in Seahorse assays.

We next conducted untargeted metabolomics on purified mouse CD8^+^ T cells to systematically characterize the global metabolic changes induced by HPK1 deletion. This global profiling identified 426 differentially regulated metabolites (Fig. S4F-G). MetaboAnalyst revealed distinctive enrichment patterns across various metabolic pathways, including the citric acid cycle, Warburg effect, carnitine synthesis, de novo triacylglycerol biosynthesis, and mitochondrial beta-oxidation of fatty acids (Fig. 4I). Given the critical role of these pathways in T cell development and fate determination^13–15^, the widespread metabolic shifts observed in HPK1-deficient cells underscore the essential role of HPK1 in regulating T cell metabolism.

Memory T cells - which are essential for long-term immune protection - require specific metabolic adaptations. HPK1-deficient CD8^+^ T cells exhibited significantly reduced levels of glycolytic intermediates (fructose-1,6-bisphosphate, glucose-1-phosphate, glucose-6-phosphate, and lactate), suggesting a metabolic shift away from glycolytic dependence (Fig. S4G-H). Concurrently, we observed an enrichment of several carnitine species, activated forms of fatty acids for mitochondrial beta-oxidation, including acylcarnitine and propionylcarnitine, in HPK1^KO^ cells (Fig. 4J-K). This metabolic signature aligns with the metabolic demands of memory T cells^39^, suggesting the enhanced reliance on fatty acid oxidation as a necessary energy source in HPK1-deficient T cells.

### Mitophagy is upregulated via HPK1 deletion

Mitophagy, a crucial form of autophagy involved in selectively removing damaged or dysfunctional mitochondria, emerges as a key regulator of memory T cell formation and maintenance^21^. Essential for mitochondrial quality control and metabolic reprogramming, mitophagy ensures optimal cellular function and metabolism. In our study, Gene Ontology (GO) analysis of RNA-seq data revealed a substantial enrichment of mitophagy-associated genes upon HPK1 deletion in mouse CD8^+^ T cells (Fig. 5A). Particularly, key mediators of mitophagy, *Pink1* and *Bnip3l*, were significantly upregulated in HPK1^KO^ cells, indicating heightened mitophagy activity in the absence of HPK1 (Fig. 5B-C). Furthermore, genes critical for autophagosome formation and activation, such as *Ulk1*, *Map1lc3a*, *Atg9b*, *Atf4*, and *Uba52*, showed increased expression in HPK1^KO^ cells, suggesting augmented autophagic processes (Fig. S5A). This enhancement in autophagy was consistently observed in human CAR T cells deficient in HPK1, as demonstrated by elevated PINK1 levels and increased autophagic flux measured via monodansylcadaverine (MDC) labeling (Fig. S5B; Fig. 5D).

**Figure 5.**
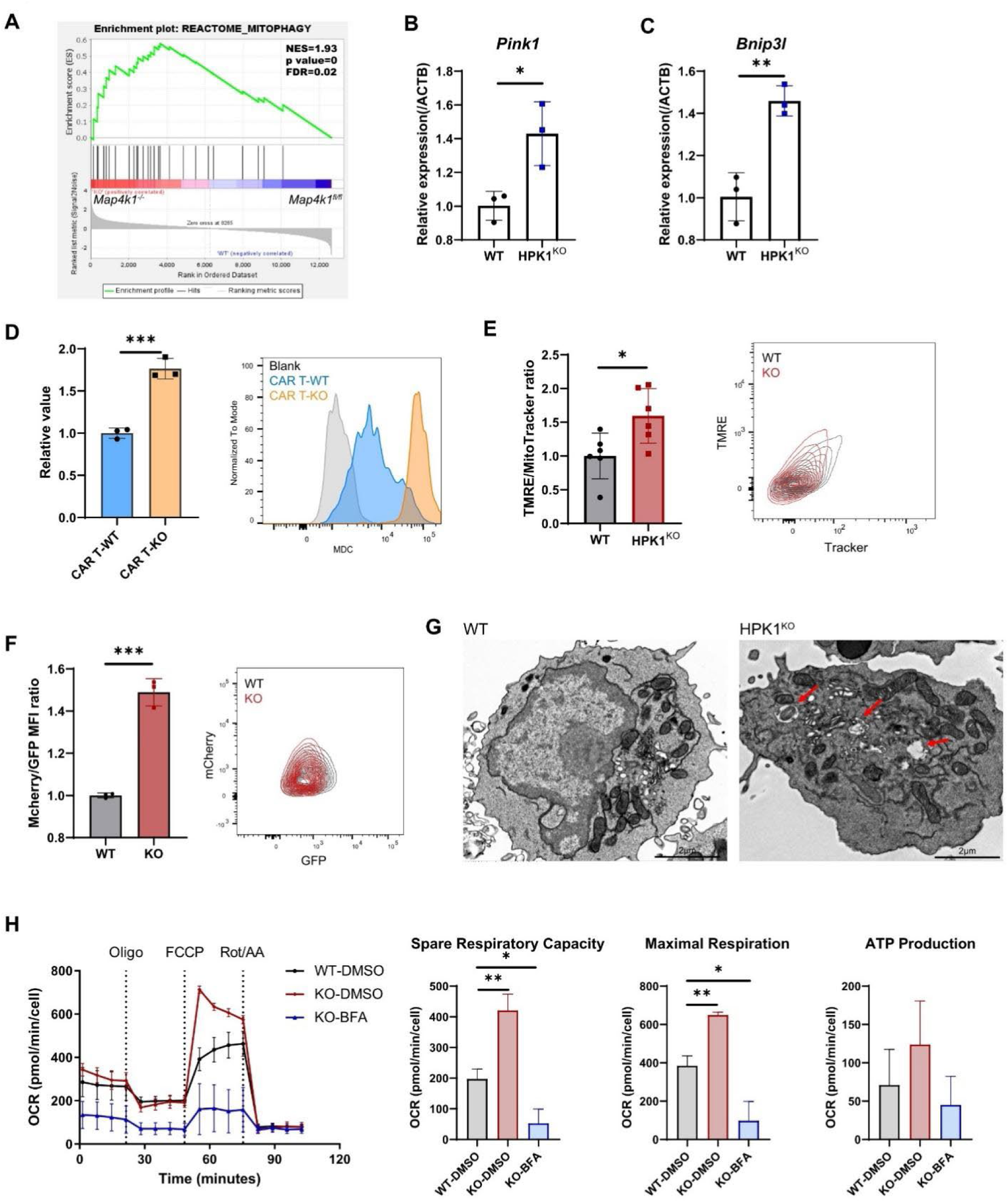
Mitophagy is upregulated due to HPK1 deletion. A. GSEA of the gene set linked to mitophagy in *Map4k1*^−/−^ versus *Map4k1*^fl/fl^ mouse CD8^+^ T cells. B-C. Expression level of *Pink1* (B) and *Bnip3l* (C) in WT and HPK1^KO^ mouse CD8^+^ T cells. D. Autophagy activity of WT and HPK1^KO^ CD19 CAR T cells measured by MDA staining. E. Mitochondrial membrane potential and mitochondrial mass of WT and HPK1^KO^ mice CD8^+^ T cells measured using TMRE and MitoTracker, respectively. The TMRE/MitoTracker ratio was calculated and normalized to WT. F. The fluorescence of mt-Keima measured in WT or HPK1^KO^ Jurkat cells via flow cytometry. G. Transmission electron micrograph of WT and HPK1^KO^ mice CD8^+^ T cells. H. OCR of mice CD8^+^ T cells treated with BFA or DMSO (left). Statistical analysis of spare respiration capacity, maximal respiration, and ATP production of indicated cells (right). Oligo, oligomycin; FCCP, carbonyl cyanide-p-trifluoromethoxyphenylhydrazon; Rot/AA, rotenone and antimycin A.

Assessment of mitochondrial function revealed a moderate increase in the ratio of mitochondrial membrane potential to mass in HPK1^KO^ cells, indicating improved mitochondrial activity per mitochondrial mass (Fig. 5E). Direct measurement of mitophagy activity using mitochondrial-targeted fluorescent protein Keima (mt-Keima) showed a significant fluorescence shift in HPK1^KO^ cells, indicative of enhanced mitophagy activity (Fig. 5F; Fig. S5C). Electron microscopy further confirmed enlarged and increased numbers of mitochondria in HPK1^KO^ cells, along with a notable rise in the proportion of cells containing autophagosomes, supporting augmented mitophagy in HPK1-deficient T cells (Fig. 5G; Fig. S5D).

These findings prompted us to investigate the functional implications of HPK1-mediated mitophagy regulation in T cell metabolism and memory formation. Given the critical role of mitochondrial turnover in maintaining metabolic homeostasis and effector functions in T cells^40^, we hypothesized that HPK1 deletion enhances mitochondrial renewal, thereby improving oxidative phosphorylation efficiency. To validate this, we pharmacologically inhibited autophagy and mitophagy using bafilomycin A1 (BFA) in mouse CD8^+^ T cells and assessed mitochondrial function with Seahorse assays. BFA treatment substantially reduced oxygen consumption rate, spare respiratory capacity, and maximal respiration in HPK1^KO^ cells (Fig. 5H), underscoring the essential role of autophagy and mitophagy in preserving mitochondrial function and metabolic balance in T cells.

### HPK1 influences mitochondrial fitness through the mTOR signaling pathway

We then aimed to elucidate the mechanisms underlying HPK1-mediated mitophagy in CD8^+^ T cells. Gene Ontology (GO) analysis of HPK1-deficient T cells revealed considerable enrichment of genes associated with ribosome biogenesis, regulation of autophagy, and the mTOR signaling pathway (Fig. 6A). Particularly, HPK1-deficient T cells exhibited upregulation of genes involved in negative regulation of mTOR signaling (Fig. 6A-B). Consistently, pseudobulk analysis of scRNA-seq data from patient-derived CD19 CAR T cells showed downregulation of ribosomal and translation-related genes in the KO group, processes tightly controlled by the mTOR pathway (Fig. S6A-B). In addition, we performed ATAC-seq sequencing of HPK1^KO^ CD19 CAR-T and WT CD19 CAR-T cells from two patients. The results showed that there were significantly more down-regulated peaks in HPK1^KO^ CD19 CAR-T cells (Fig. S6C), indicating that HPK1 knockdown affects chromatin openness in CAR-T cells. Most of the differential peaks were concentrated in the promoter region of the genome (Fig. S6D), suggesting that these differential peaks were closely linked to gene functions. Moreover, the peak in the promoter region of the mTOR gene was significantly down-regulated in the KO group (Fig. 6C). The chromatin accessibility of genes within the mTOR pathway were obviously reduced in HPK1-depleted CAR T cells (Fig. S6E). These findings highlight the intricate regulatory role of HPK1 in shaping chromatin landscape and gene expression patterns associated with mTOR signaling, influencing downstream cellular processes such as autophagy and mitochondrial dynamics.

**Figure 6.**
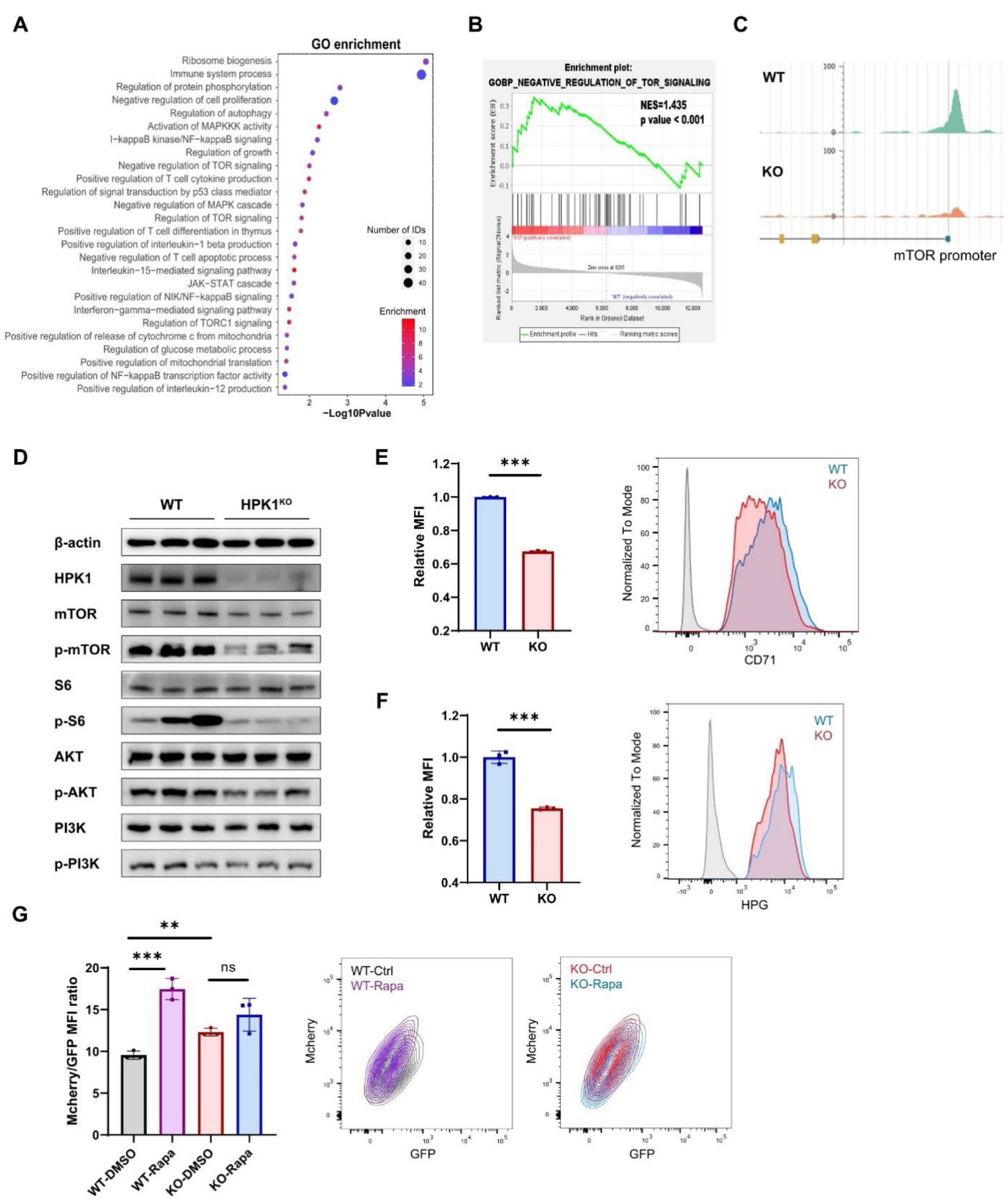
HPK1 influences mitochondrial fitness through the mTOR signaling pathway. A. Gene Ontology (GO) enrichment of RNA-seq data of *Map4k1*^−/−^ versus *Map4k1*^fl/fl^ mouse CD8^+^ T cells. B. GSEA of TOR signaling pathway in *Map4k1*^−/−^ versus *Map4k1*^fl/fl^ mouse CD8^+^ T cells. C. The peak in the promoter region of mTOR gene from ATAC-seq results using patient-derived samples. D. Immunoblot of indicated proteins of WT and HPK1^KO^ mouse CD8^+^ T cells. E. Expression of CD71 in WT and HPK1^KO^ CAR T cells detected by FACS. F. Rate of protein synthesis measured by HPG incorporation. G. Fluorescence of mt-Keima in WT or HPK1^KO^ Jurkat cells treated with rapamycin or DMSO.

Further examination of the mTOR pathway in HPK1-deficient T cells revealed alterations in key signaling nodes. CD8^+^ T cells from HPK1^KO^ mice displayed significantly reduced phosphorylation levels of AKT and mTOR proteins (Fig. 6D). Consistent with this, phosphorylation of S6 protein, a downstream target of mTORC1, was dramatically decreased in *Map4k1^−/−^* cells (Fig. 6D). Flow cytometry analysis showed decreased CD71 expression (Fig. 6E; Fig. S6F), a marker indicative of mTORC1 pathway activity, further confirming the mTOR pathway downregulation in HPK1-deficient T cells. Moreover, to assess mTOR pathway function in protein synthesis, we observed slower protein synthesis in *Map4k1^−/−^* cells using an HPG labeling assay (Fig. 6F).

To explore the functional implications of mTORC1 modulation on HPK1-associated mitophagy, we pharmacologically targeted the mTOR pathway using rapamycin, a potent mTORC1 inhibitor (Fig. 6G; Fig. S6G-H). Remarkably, rapamycin treatment significantly enhanced mitophagy in WT CD8^+^ T cells, evidenced by a notable shift in mt-Keima fluorescence. In contrast, rapamycin had minimal effect on mitochondrial activity in HPK1^KO^ cells, suggesting that HPK1 depletion may render T cells less responsive to mTORC1 inhibition. This observation underscores the complex interplay between HPK1 and the mTOR pathway in regulating mitochondrial function and cellular metabolism.

## Discussion

Enhancing the formation of memory CD8^+^ T cells, which retain plasticity and sustain T cell stemness and robust recall function, holds promise for advancing immunotherapy. Despite its recognized role in regulating T cell immune responses, the influence of HPK1 on T cell memory differentiation has remained unclear. Here, we demonstrate that HPK1 downregulation promotes preferential differentiation of CD8^+^ T cells towards a memory phenotype through metabolic reprogramming. Both acute infection and cancer models in *Map4k1*^−/−^ mice show increased proportions of CD62L^+^CD27^+^ memory CD8^+^ T cells, accompanied by enhanced proliferative capacity, effector function, and prolonged antitumor efficacy. Validation with patient-derived samples underscores the clinical relevance of these findings. The enhanced memory differentiation is likely attributed to improved mitochondrial quality through elevated mitophagy and oxidative metabolism in HPK1-deficient cells. This study marks the first evidence that HPK1 expression negatively correlates with memory CD8^+^ T cell formation.

Given HPK1’s broad impact beyond CD8^+^ T cells, it is plausible that HPK1 influences memory subtype formation in other immune cells. Both CD4^+^ T cells and CD8^+^ T cells are governed by TCR signaling, which HPK1 negatively regulates akin to B cell receptor (BCR) signaling through similar molecular processes^41^. Further exploration is needed to understand these effects comprehensively on CD4^+^ T cells and B cells. HPK1 also restrains NK cell activation, pivotal in innate immune responses against tumors^35^, despite suggestions of NK cell memory plasticity^42–44^. This prompts investigation into whether HPK1 affects NK cell stemness. Given HPK1’s influence via signaling pathways such as NF-κB^45,46^ and MAPK^28,29,47^, which impact differentiation, survival, and effector functions across immune cell subsets, studying HPK1’s role in other immune cells promises insights into their coordinated responses to re-encountered challenges.

In addition to affecting these pathways, we found that HPK1 expression negatively correlates with genes related to mTOR signaling. mTOR signaling activation dynamically changes upon TCR stimulation and closely influences T cell differentiation^48–50^. Inhibition of mTOR by rapamycin during acute LCMV infection enhances memory T cell formation^26^ and improves immunotherapy efficacy^51,52^. HPK1 depletion reduces mTOR phosphorylation and downstream targets like CD71 and p-S6, promoting memory differentiation. Paradoxically, while TCR activation increases mTOR signaling, HPK1 depletion shows decreased phosphorylation of mTOR and S6 30 days post LCMV-Armstrong infection, suggesting HPK1 exerts stage-dependent effects on the mTOR pathway. Further studies are underway to elucidate these mechanisms.

Moreover, HPK1-deficient cells exhibit enriched gene sets associated with metabolism changes, potentially linked to HPK1 kinase activity and MAPK pathway alterations. Metabolites crucial for epigenetic modification, such as S-adenosylmethionine (SAM) and S-adenosylhomocysteine (SAH), are altered in HPK1-deficient cells, suggesting a link between metabolism and transcriptional changes. Notably, HPK1 deficiency enhances mitochondrial metabolism, boosting oxidative phosphorylation and fatty acid oxidation. By inhibiting the mTOR pathway, HPK1 regulates mitophagy dynamics, crucial for maintaining mitochondrial quality and metabolic fitness essential for memory T cell formation. These metabolic shifts align with the observed memory T cell phenotype, highlighting HPK1’s role in enhancing T cell metabolic fitness to sustain effector functions and promote memory formation. Given the critical role of metabolic reprogramming and mitochondrial fitness in T cell fate decisions, manipulating metabolic enzyme activities like proline dehydrogenase 2 (PRODH2), mitochondrial pyruvate carrier (MPC), and acylglycerol kinase (AGK) can influence memory cell formation^18,53,54^. The activity of mitophagy is critical to preserving mitochondrial quality and metabolic fitness necessary for memory T cell formation^21,55,56^. Our findings underscore the cross-talk among HPK1, metabolism, and T cell fate, advocating for targeting HPK1-mediated signaling or metabolic pathways to enhance memory T cell responses.

## Competing Interests

The authors declare that they have no conflict of interest.

## Acknowledgments

We are grateful to Xi’an Yufan company for CAR T cells construction and amplification. We thank Chenguang Zhao, Pengcheng Jiao, and Jiaojiao Ji from the Center of Biomedical Analysis of Tsinghua University for helping in TEM transmission electron microscopy sample preparation and flow cytometry analysis. We thank Lina Xu from the Protein Research and Technology Center of Tsinghua University for helping with metabolomics analysis.

## Author contributions

X.L. and L.Y. designed the research. L.Y. and X.G performed most of the wet-lab experiments. T.W. performed most of the bioinformatics analysis. C.H., H.S. and H.W. helped with the mouse experiment. L.Y. and T.W. wrote the manuscript with the help from X.L., all authors reviewed and edited the manuscript.

## Materials and Methods

### Mice

Map4k1^fl/fl^ C57BL/6J mice and CD8^Cre^ C57BL/6J mice were obtained from Cyagen Biosciences. HPK1^KO^ C57BL/6J mice, OT1 mice, and CD45.1^+^ congenic mice were bred and maintained in our lab. Map4k1^fl/fl^ C57BL/6J mice were crossed with CD8^Cre^ C57BL/6J mice to generate mice with specific HPK1 depletion in CD8^+^ cells (referred to as *Map4k1*^−/−^ mice). Mice aged between 6 to 15 weeks were used for all experiments. All animal procedures were conducted in accordance with the guidelines of the Laboratory Animal Research Center of Tsinghua University and approved by the International Association for Assessment and Accreditation of Laboratory Animal Care.

### Infection models

Lymphocytic choriomeningitis virus Armstrong strain (LCMV-Arm) was provided by Dr. Chen Dong’s lab, propagated in-house, and used for intraperitoneal (i.p.) infection of mice with 2×10^5^ plaque-forming units (pfu). Mice were sacrificed on day 8, day 20, and day 30 post-infection for flow cytometry analysis.

### Cell proliferation assay

CD8^+^ cells isolated from mice at day 30 post-LCMV-Arm infection were labeled with 5 μM CellTrace ™ CFSE (Invitrogen) and cultured in a medium containing 1 μg/mL gp33 peptide and 10 ng/mL IL-2 for four days. CFSE dilution was analyzed by flow cytometry.

### Tumor models

A total of 1×10^5^ B16-OVA cells were subcutaneously injected into Map4k1^fl/fl^ mice or Map4k1^−/−^ mice, and tumor growth was monitored every three days. In adoptive transfer experiments, naïve WT or HPK1^KO^ mice were vaccinated twice with 100 μg OVA peptides mixed with complete Freund’s adjuvant (1:1). A total of 1×10^6^ CD8^+^ cells sorted from vaccinated mice were transferred intravenously into naïve CD45.1^+^ recipient mice, followed by injection of 5×10^5^ B16-OVA cells subcutaneously. Tumor volume was calculated using the formula (length×width^2^)/2, and mice were euthanized when tumors reached 2,000 mm^3^.

### Recall assays

For *in vivo* recall assays, total CD8^+^ cells isolated from spleens of infected mice using MojoSort™ Mouse CD8 T Cell Isolation Kit (Biolegend) (1×10^5^) were transferred intravenously into naïve CD45.1^+^ mice and re-challenged with 2×10^5^ pfu of LCMV-Arm or 100 μg OVA-peptides mixed with complete Freund’s adjuvant. Results were analyzed at day 6 or day 3 post-re-challenge.

For *in vitro* recall assays, CD8^+^ cells isolated from infected mice were stimulated with 1 μg/mL gp33 peptides in the presence of Brefeldin A (Golgiplug, BD Bioscience) for 4 hours prior to flow cytometry analysis.

### Oxygen Consumption Rate

OCR was measured using a Seahorse XF9 analyzer (Agilent). CD8^+^ cells activated with anti-CD3 antibody (2 mg/mL, coating plate) and anti-CD28 antibody (1 mg/mL) for 12 hours were pretreated with non-buffered XF medium (non-buffered RPMI 1640 containing 10 mM glucose, 1 mM sodium pyruvate, and 2 mM glutamine). Cells were seeded at 2×10^5^ cells per well in XF96 cell culture microplates coated with poly-lysine. The cells were incubated in a non-CO_2_ incubator for 1 hour at 37 °C. Sequential injections of 1.25 μM oligomycin, 50 mM 2-deoxy-d-glucose, 1.5 μM FCCP, 1 μM rotenone, and 1 μM antimycin A were performed at indicated times and OCR was measured.

### Flow cytometry

Cells from spleens or lymph nodes were isolated by grinding spleens through 70 μm filters, treated with red blood cell lysis solution (Solarbio), and washed with PBS. Cells were stained with antibodies in PBS containing 3% FBS for 30 minutes at room temperature. To detect Granzyme B and cytokine production, cells were stimulated *in vitro* using 1 μg/mL gp33 peptides in the presence of Brefeldin A (Golgiplug, BD Bioscience) for 4 hours, followed by fixation and permeabilization using a fixation/permeabilization kit (BD Biosciences).

### RNA isolation, RNA-seq, and qPCR

Splenic CD8^+^ cells were isolated using a MojoSort™ Mouse CD8 T Cell Isolation Kit (Biolegend) following the manufacturer’s instructions. Approximately 1×10^6^ cells per sample (n=3, each sample mixed 3 mice together) were utilized for total RNA extraction using Trizol (Solarbio). RNA-seq libraries were constructed and sequenced by GENEWIZ. For qPCR analysis, 1 μg of total RNA was reverse transcribed (Vazyme), and 20 ng cDNA was used per reaction using specific PCR primers.

### Metabolomic analysis

A total of 2×10^7^ splenic CD8^+^ cells per sample (n=3, each sample mixed 3 mice together) were washed with pre-cooled PBS twice and resuspended in 80% cold methanol by vigorous vortexing. Samples were extracted at −80 °C overnight and centrifuged at 12,000 × *g* for 10 minutes at 4 °C. The supernatant was obtained, and the extracts were detected using LC-MS/MS. Raw data were processed using Compound Discoverer (v3.1, Thermo Fisher) and MetaboAnalyst was used for subsequent data analysis.

### Mitochondrial mass and membrane potential analysis

Mitochondrial mass and membrane potential were examined using MitoTracker Deep Red FM (YEASEN) and tetramethylrhodamine ethyl ester (TMRE, Beyotime). Cells were stained with 50 nM MitoTracker Deep Red FM and TMRE (1×, diluted following instruction) in a medium lacking FBS at 37 °C (5% CO_2_) for 30 minutes. Cells were rinsed with PBS, stained, and analyzed by flow cytometry.

### Mitochondrial respiration complex activity analysis

Freshly sorted CD8^+^ cells (n=3, each sample mixed 3 mice together) from OVA-vaccinated mice were characterized using CheKine^TM^ mitochondrial respiration complex activity kits (Abbkine). Cells were rinsed with cold PBS and incubated in Reagent I with protease inhibitor for 30 minutes on ice. The cells were lysed using a 22-G needle, blowing about 25 times, followed by centrifugation at 600 × *g*, 4 °C for 5 minutes. The supernatant was collected and centrifuged again at 12,000 × *g*, 4 °C for 15 minutes. The precipitates containing extracted mitochondria were characterized following the manufacturer’s instructions.

### Mitochondrial morphology analysis

A total of 1×10^7^ splenic CD8^+^ cells per sample (n=3, each sample mixed 3 mice together) from OVA-vaccinated mice were immediately fixed using a fixation buffer (2.5% glutaraldehyde, 0.1 M phosphate buffer [PB], pH 7.4) for 1 hour at room temperature and overnight at 4 °C. Mitochondrial morphology was characterized using Hitachi 7650B transmission electron microscopy.

### MDC assay

Monodansylcadaverine (MDC) staining was utilized to detect autophagy activity following the manufacturer’s instructions (Beyotime). Human CAR T cells cultured for 30 days *in vitro* were activated using anti-CD3 and anti-CD28 antibodies for 48 hours. Cells were rinsed with PBS and stained using diluted MDC for 30 minutes at 37 °C. After being washed three times with assay buffer, cells were analyzed by flow cytometry with a 405 nm laser.

### Statistical analysis

Results are presented as means ± standard deviation. However, for the tumor growth curve, means ± standard error was used. Statistical analyses were performed by using GraphPad Prism 8. A two-tailed Student’s *t-*test was used to analyze the differences between groups. ANOVA was used to analyze two individual response groups. The significance level was established at \**P*-value < 0.05, \*\**P*-value < 0.01, \*\*\**P*-value < 0.001, \*\*\*\**P*-value < 0.0001, n.s. not significant.

---

**Figure S1.**
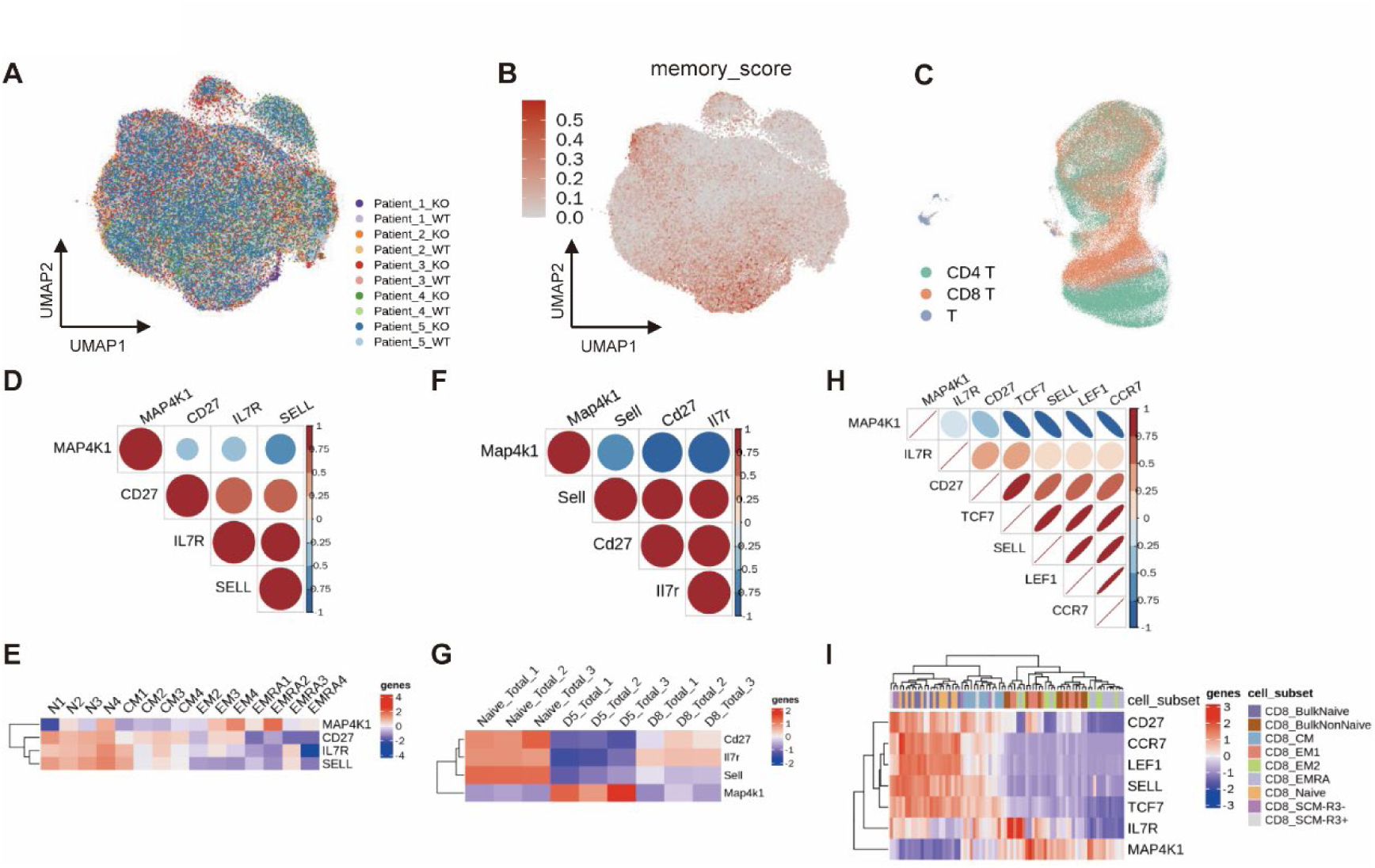
*MAP4K1* expression is negatively correlated with *SELL* and *CD27* in human and mouse T cells. A. Uniform Manifold Approximation and Projection (UMAP) visualization of single-cell RNA sequencing data shows the distribution and clustering of CAR-T cells across experimental conditions, with color-coding representing distinct samples; B. The distribution of memory_score in CAR-T cells; C. The distribution of cell types in dataset of Deng et al., colors represent distinct cell types; D-E. *MAP4K1* expression shows inverse correlations with memory-associated markers *SELL* and *CD27* in Willinger’ human CD8^+^ T cell microarray dataset. D. gene expression correlation plot, circle color and size represent the correlation values; E. Heatmap of relevant gene expression across different CD8^+^ T cell types. Colors indicate gene expression levels. F-G. *MAP4K1* expression exhibits inverse correlations with *SELL* and *CD27* in Araki’ mouse T cell microarray dataset. F. gene expression correlation plot, circle color and size represent the correlation values; G. Heatmap of relevant gene expression across different T cell types. Colors indicate gene expression levels. H-I. *MAP4K1* expression shows inverse correlation with *SELL* and *CD27* in Giles’ human CD8^+^ T cell RNA-seq dataset. H. gene expression correlation plot, circle color and size represent the correlation values; I. Heatmap of relevant gene expression across different CD8^+^ T cell types. Colors indicate gene expression levels.

**Figure S2.**
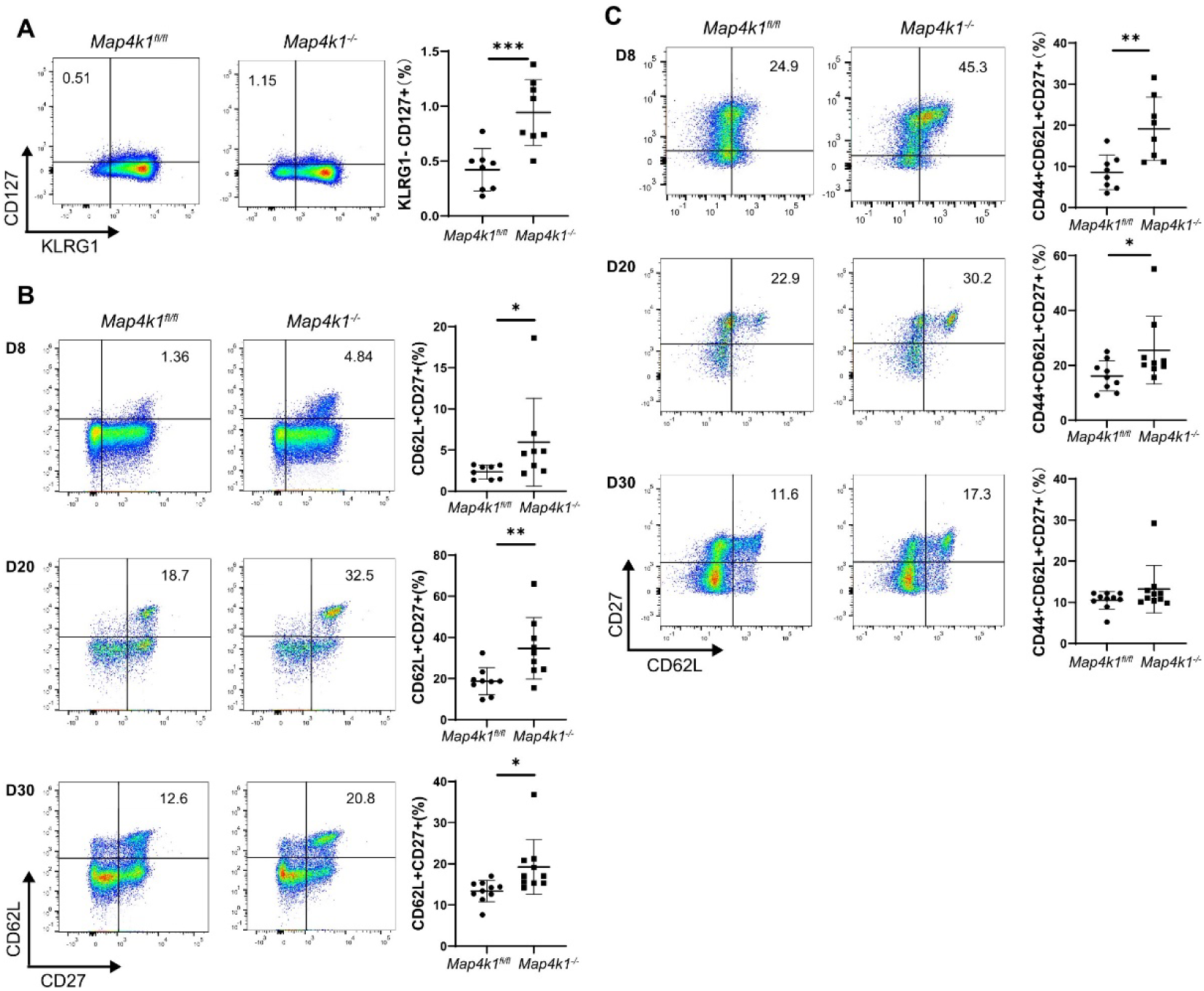
HPK1-defect mice favor memory CD8^+^ T cell differentiation. A. Flow cytometry (left) and quantification of the frequency (right) of CD127^+^KLRG1^−^cells in total CD8^+^ T cells from lymph node at day8 after infection. B-C. Flow cytometry (left) and quantification of the frequency (right) of CD62L^+^CD27^+^ cells (B) and CD44^+^CD62L^+^CD27^+^ cells (C) in total CD8^+^ T cells from lymph node at indicated time after infection.

**Figure S3.**
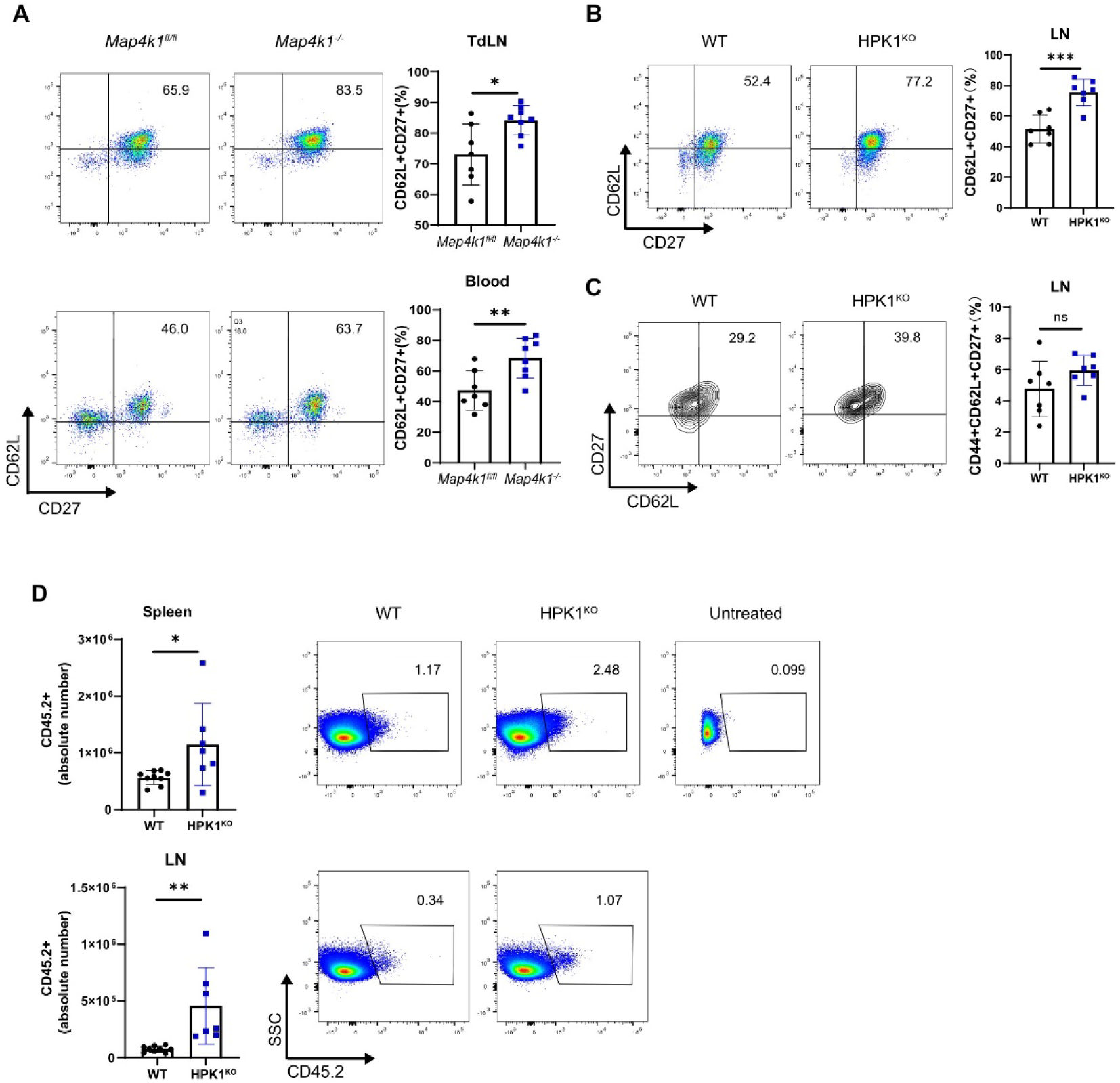
Depletion of HPK1 in CD8^+^ cell improves long-term antitumor capacity. A. FACS analyses of CD62L and CD27 expression of CD8^+^ T cells from tumor-draining lymph nodes at day15. B-C. Flow cytometry (left) and quantification of the frequency (right) of CD62L^+^CD27^+^ cells (B) and CD44^+^CD62L^+^CD27^+^ cells (C) in total CD8^+^ T cells of lymph nodes at day28 after OVA vaccination. D.FACS analyses of donor-derived CD8^+^ cells in spleen (upper) and lymph node (bottom) after the OVA peptide re-challenge.

**Figure S4.**
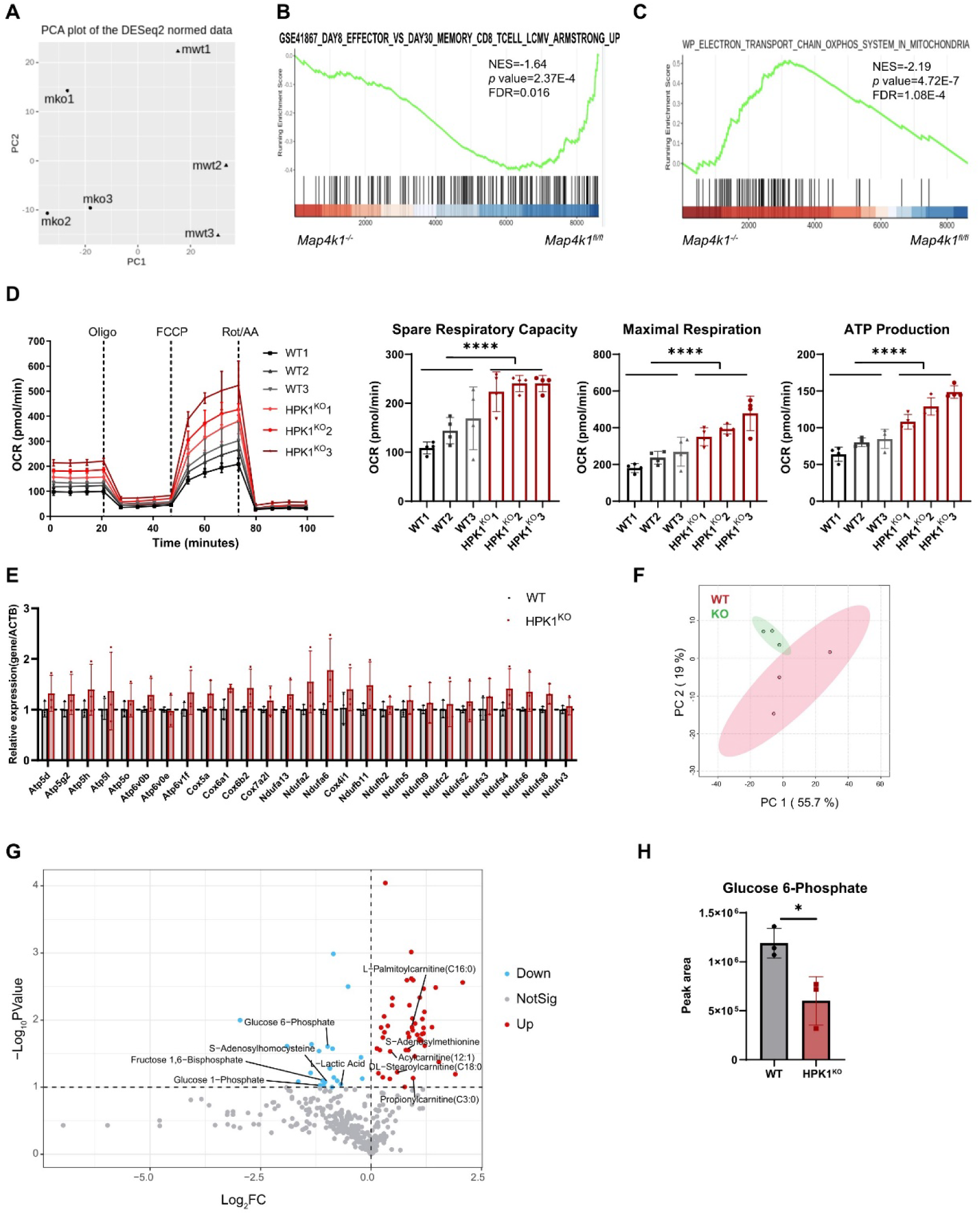
HPK1 influences CD8^+^ T cell metabolic reprograming. A. Principal component analysis (PCA) of RNA-seq data from *Map4k1*^fl/fl^ and *Map4k1*^−/−^ mice CD8^+^ T cells. B-C. GSEA plots of indicated gene sets in *Map4k1*^−/−^ compared to *Map4k1*^fl/fl^ mice CD8^+^ T cells. D. OCR of WT and HPK1^KO^ mice CD8^+^ T cells. (left). Statistical analysis of spare respiration capacity, maximal respiration and ATP production of indicated cells (right). Oligo, oligomycin; FCCP, carbonyl cyanide-p-trifluoromethoxyphenylhydrazon; Rot/AA, rotenone and antimycin A. E. Relative gene expression of indicated genes of WT and HPK1^KO^ mice CD8^+^ T cells measured by quantification PCR. F. PCA plot of metabolomics data of WT and HPK1^KO^ mice CD8^+^ T cells. G. Volcano plot of differentially represented metabolites compared WT and HPK1^KO^ mice CD8^+^ T cells. Blue dots indicate decreased metabolites, red dots indicate increased metabolites and grey dots indicate not-significantly changed metabolites. H. Quantification of glucose-6-phosphate by LC-MS in CD8^+^ T cells from WT and HPK1^KO^ mice.

**Figure S5.**
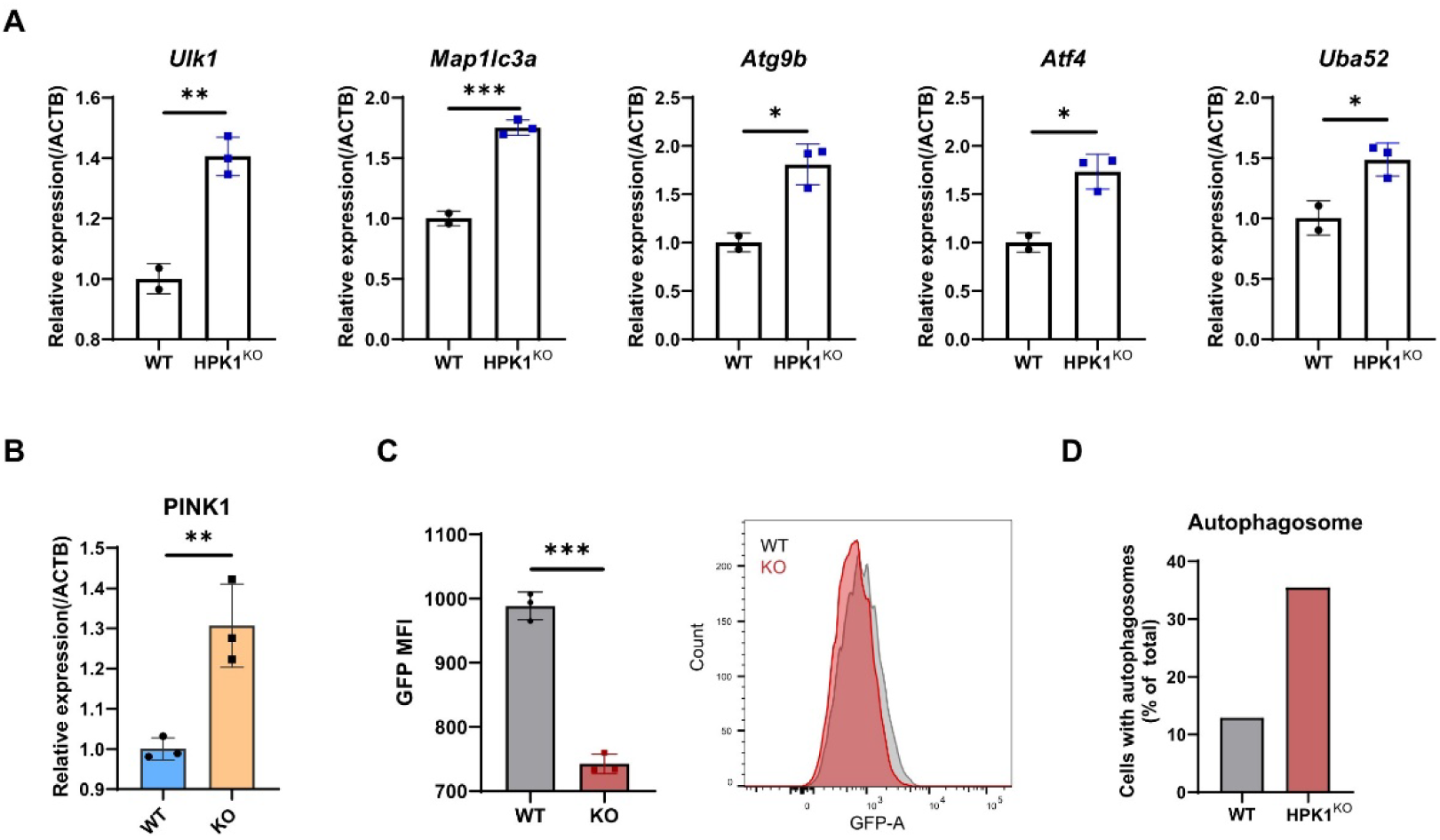
Mitophagy is up-regulated due to HPK1 deletion. A-B. qPCR measured the expression of indicated genes in WT and HPK1^KO^ mice CD8^+^ T cells (A) or human CAR T cells (B). C. The intensity of GFP fluorescence of mt-Keima protein in WT and HPK1^KO^ Jurkat cells. D. The percentage of cells containing autophagosomes in transmission electron microscope graph of WT and HPK1^KO^ mice CD8^+^ T cells.

**Figure S6.**
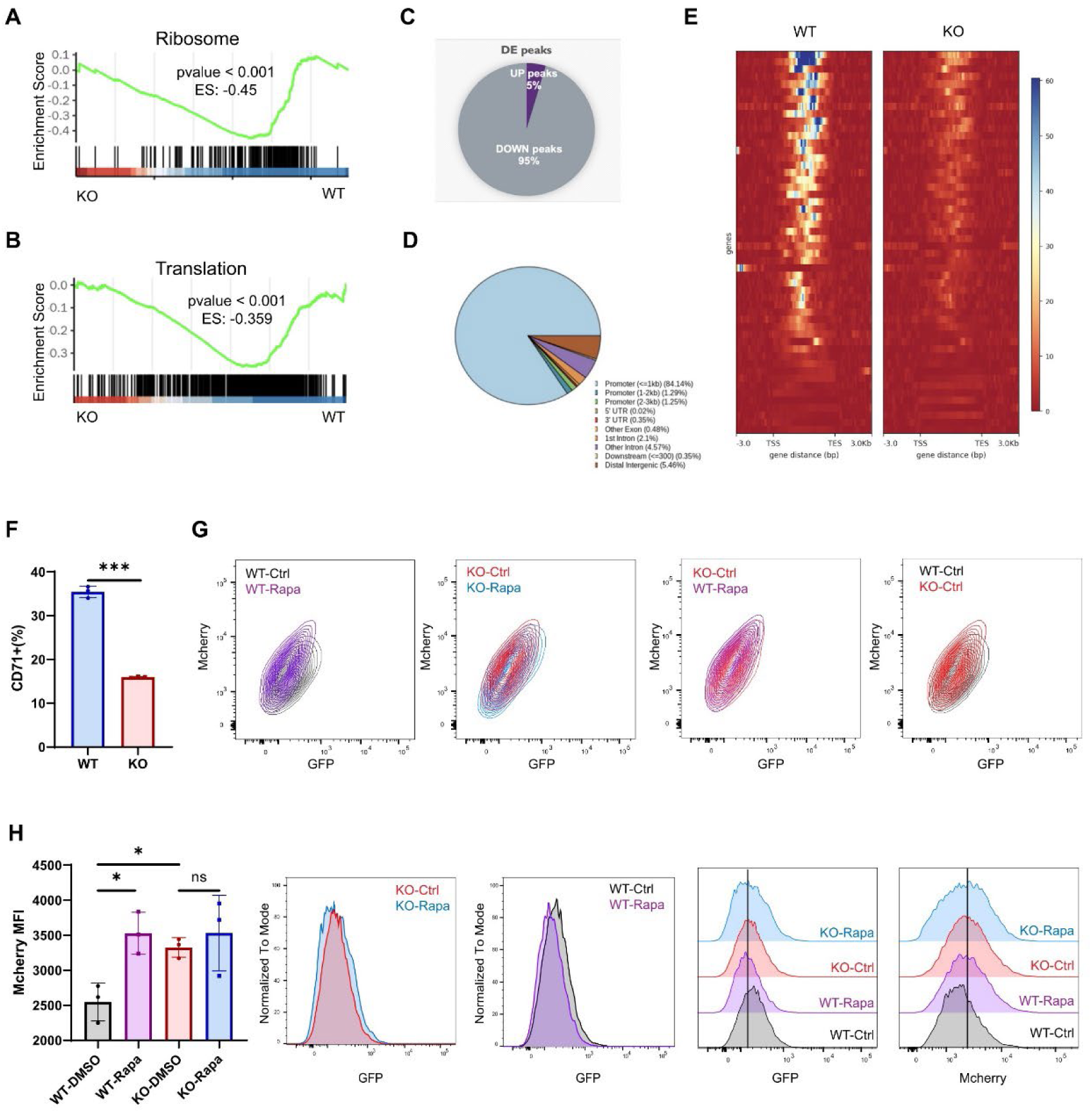
HPK1 influences mitochondrial fitness through mTOR signaling pathway. A-B. GSEA plots of indicated gene sets enriched in HPK1^KO^ compared to WT patient-derived CD19 CAR-T cells. C-E. Results of ATAC-seq of HPK1^KO^ and WT CD19 CAR-T cells from two patients. Statistical analysis (C) and gene location (D) of differential peaks. Heatmap of ATAC-seq peaks of genes within mTOR pathway (E). F. The percentage of CD71^+^ cells in WT and HPK1^KO^ mice CD8^+^ T cells. G-H. Flow cytometry plot (G) and the intensity of GFP or mcherry fluorescence (H) of mt-Keima in WT or HPK1^KO^ Jurkat cells treated with rapamycin or DMSO.

## Notes

### Competing Interest Statement

The authors have declared no competing interest.

